# Binding of dsRNA by *D. melanogaster* Dicer-2 is substrate-dependent and regulated by Loquacious-PD

**DOI:** 10.1101/2020.11.19.390161

**Authors:** M. Jonely, R. K. Singh, B. L. Bass, R. Noriega

## Abstract

*Drosophila melanogaster* Dicer-2 is a large, multidomain protein that cleaves double-stranded RNA (dsRNA) into small interfering RNAs in a terminus-dependent manner as part of the RNA interference pathway. We characterize the local binding environment involved in this substrate-selective molecular recognition event by monitoring the time-resolved photophysics of a cyanine dye linked to the dsRNA terminus. We observe substantial changes in the molecular rigidity and local freedom of motion of the probe as a function of distinct conformations of the biomolecular complex between Dicer-2 and dsRNA as a function of dsRNA termini, the presence of regulatory proteins, and the addition of a biochemical energy source (ATP) or a non-hydrolysable equivalent (ATP-γS). With a clustering analysis based solely on these molecular-scale measures of the local binding environment at the dsRNA terminus, we identify sub-populations of similar conformations that define distinct modes of molecular recognition which are correlated with biochemical activity. These observations reveal the important role of substrate-selective molecular recognition properties for proteins with multiple domains that can bind RNA, regulatory proteins, and cofactors.

**STATEMENT OF SIGNIFICANCE:** The molecular-scale determinants of protein-RNA binding remain elusive, particularly when different subunits of a single protein confer specificity toward small structural differences of their RNA partners. An important case is that of *Drosophila melanogaster* Dicer-2, a critical component of the antiviral RNA interference response. Dicer-2 discriminates between double stranded RNA with blunt or 3’ overhang termini, a feature suggested to mediate recognition of “self” vs. “non-self” substrates. We study these interactions at the binding site with a fluorescent label at the RNA terminus, monitoring intramolecular and collective measures of flexibility to report on the local environment. Dicer-2 has distinct modes of molecular recognition which are regulated by accessory proteins and ATP, leading to different conformations and tuning biochemical activity.

## INTRODUCTION

Molecular recognition and binding between large biomolecules are critical components to many biological processes; both in healthy cellular function and in the course of disease.(1–4) Our understanding of these interactions relies on a combination of structural biology tools and biochemical studies.(5–13) However, the role of the local binding environment in biomolecular recognition remains to be fully understood in protein-RNA interactions. Difficulties in the characterization of molecular recognition in biomolecular complexes stem primarily from the large number of conformational degrees of freedom of the individual macromolecular components, the substantial structural changes that take place over the course of initial binding and as a result of subsequent biological function, and the manifold interactions among protein residues, nucleobases, solvent molecules, ions, and small molecule cofactors.(14–16) Importantly, these local environments are far from static – they experience multiscale dynamics ranging from fast short-range motions (ps–ns, nm) to much slower long-range events (ms, μm).(17–21) In order to understand the relationship between local binding environment and the mode of molecular recognition in protein-RNA complexes, it is crucial to interrogate such local environments from a molecular perspective that includes their dynamic behavior.

Biological systems have evolved to perform complex molecular recognition tasks in crowded environments with exquisite specificity. A particularly interesting feature of molecular recognition in biology is the ability of some proteins to tune their substrate specificity through the incorporation of multiple binding domains.(22–24) One of such proteins is *D. melanogaster* Dicer-2 (dmDcr-2; PDB: 6BUA, 6BU9), a necessary component of the antiviral RNA interference pathway that cleaves double-stranded RNA (dsRNA) intermediates into small interfering RNA (siRNA) products.(25, 26) Dicer-2 is a large enzyme (1724 amino acid residues, ~198 kDa) with multiple sub-domains (**Fig. 1**) that play distinct roles in its substrate-specific processing of blunt-end dsRNA substrates (BLT) vs. those with a 2-nucleotide overhang at the 3’ terminus (3’ovr).(24, 27, 28) Important features of Dicer-2 function are summarized below.

**Figure 1.**
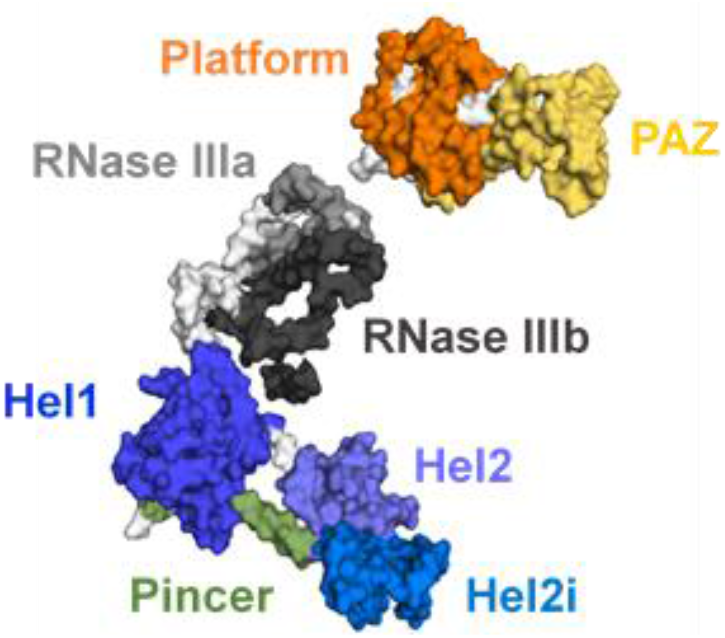
Structure of apo dmDcr-2 (PDB: 6BUA) where its various domains are identified: tandem RNase III domains (RNase IIIa and RNase IIIb), three helicase subdomains (Hel1, Hel2, and Hel2i), a pincer, a platform, and a PAZ (Piwi-Argonaute-Zwille) domain.

### Binding site

BLT dsRNA interacts with the helicase domain of dmDcr-2, while 3’ovr dsRNA engages the platform•PAZ domain at the opposite end of the molecule (**Fig. 1**).(24) We note that recent transient kinetic studies indicate some overlap of binding sites;(29) however, in this study we focus on equilibrium conditions.

### ATP dependence

Consistent with the clear role of ATP to helicase functions,(30) ATP significantly improves binding of BLT dsRNA to dmDcr-2’s helicase domain (e.g., for 52 basepair BLT dsRNA 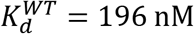 without ATP, 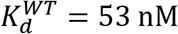 with ATP), and ATP hydrolysis is required for cleavage of BLT dsRNA. On the contrary, ATP is not required to cleave 3’ovr dsRNA, and it decreases the affinity of dmDcr-2 for 3’ovr dsRNA (e.g., for 52 basepair 3’ovr dsRNA 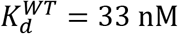 without ATP, 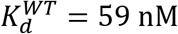 with ATP).(27, 28)

### Reactivity

BLT dsRNA is cleaved processively, with multiple siRNAs produced prior to dissociation. Current models indicate dsRNA bound at the helicase domain is threaded through the helicase domain to encounter the RNase III nuclease domains, and correspondingly, single-turnover experiments show that the reaction of BLT dsRNA yields products that are heterogeneous in length. By contrast, cleavage of 3’ovr dsRNA is distributive and leads to siRNAs with a well-defined size (~22-nt).(24)

### Cleavage rate

BLT substrates are cleaved at a substantially faster rate than 3’ovr substrates (e.g., 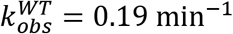 for 106 basepair BLT substrates vs. 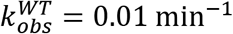 for 106 basepair 3’ovr substrates).(27)

### Regulatory proteins

The activity of dmDcr-2 is modulated by accessory factors. In particular, its partner protein Loquacious-PD (Loqs-PD) modulates dmDcr-2’s termini-dependent activity (e.g., in the presence of Loqs-PD and ATP, the cleavage rate for 106 basepair 3’ovr dsRNA is only ~2X slower than the cleavage rate for 106 basepair BLT dsRNA).(27, 31)

The biochemical properties of dmDcr-2, especially its use of multiple domains to discriminate dsRNA substrates based on their termini, have been proposed as a mechanism to differentiate between “self” and “non-self” dsRNA as it evolved its function within the RNA interference antiviral response.(24, 32, 33) Importantly, such complex biochemical activity relies on effective, substrate-specific molecular recognition.

In this study, we identify three distinct modes of molecular recognition of dsRNA by Dicer-2 whose interactions are mediated by multiple protein domains and are strongly dependent on the presence of regulatory proteins and nucleotide-induced conformational changes: two substrate-dependent types of interaction in the absence of Loqs-PD, and a substrate-independent type of binding in the presence of Loqs-PD. To probe the local binding environments, we measure the polarization-resolved ultrafast fluorescence of chromophores covalently bound to the dsRNA terminus to probe the local environment at the binding site. The photophysical behavior of these fluorescent probes – fluorescence lifetime and ultrafast fluorescence anisotropy decay – report on the probe’s ability to undergo intramolecular conformational changes and participate in collective reorientations involving the surrounding solvent molecules. Because multiple protein domains are involved in the function of dmDcr-2, we interrogate their contributions to the molecular recognition process by comparing the wild type protein (dmDcr-2^WT^) with several variants in which relevant domains have been altered: deletion of helicase and pincer domains (dmDcr-2^ΔHel^), removal of RNA cleaving ability (dmDcr-2^RIII^), and alteration of platform•PAZ binding (dmDcr-2^PP^). We characterize the local binding environment for BLT and 3’ovr substrates as a function of the presence of a hydrolysable chemical energy source (ATP) or a non-hydrolysable equivalent (ATP-γS), as well as the Loqs-PD regulatory protein. We find that the local environment sensed at the RNA substrate terminus is characteristic of its binding mode and can be used as a proxy for molecular recognition.

## EXPERIMENTAL

### Materials and Methods

Protein and RNA materials used in this work were obtained and employed as detailed in previous reports.(24, 27–29, 31)

#### Proteins

Besides the wild type (dmDcr-2^WT^), several variants were used in this work: a mutant whose helicase and pincer domains have been deleted entirely (dmDcr-2^ΔHel^), a variant with a single point mutation in each RNase III domain to prevent cleavage (dmDcr-2^RIII^), and a variant with 5 point mutations in the platform•PAZ domain to alter substrate binding (dmDcr-2^PP^, originally obtained from R. Fukunaga).(24, 27, 28, 34) While the detailed biochemical activity of dmDcr-2^PP^ is not fully known, it is included here to complement observations in the remaining variants. Full-length Loqs-PD was purified and used in a manner consistent with previous reports.(27, 31) Details on the sequence modifications for protein variants used and their expression and purification are provided in the Supporting Information.

#### Nucleic acids

Sense RNA strands with a Cy3 fluorescent label at the 5’ end were purchased from Integrated DNA Technologies. Antisense RNA strands were chemically synthesized following previously reported protocols,(24) using reagents from Glen Research (Sterling VA) and HPLC purified at the DNA/Peptide Facility, part of the Health Science Center Cores at the University of Utah. Single strands were purified after polyacrylamide gel electrophoresis (17% denaturing). Directional binding of dsRNA by dmDcr-2 was facilitated by including two deoxynucleotides at the 3’ end of sense strands, and a biotin at the 5’ end of antisense strands. Sequences for sense and antisense strands are provided below:

<u>52-nt Sense strand</u>

5’-Cy3-GGAGGUAGUAGGUUGUAUAGUAGUAAGACCAGAC CCUAGACCAAUUCAUGCC-3’

*<u>CC</u>: deoxynucleotides

<u>52-nt BLT Antisense strand</u>

Biotin-5’-GGCAUGAAUUGGUCUAGGGUCUGGUCUUACUACUAUACAACCUACUACCUCC-3’

<u>54-nt 3’ovr Antisense strand</u>

Biotin-5’-GGCAUGAAUUGGUCUAGGGUCUGGUCUUACUACUAUACAACCUACUACCUCCAA-3’

To prepare dsRNAs from the single RNA strands above, equimolar amounts of 52-nt sense strands and either 52-nt or 54-nt antisense strands (for BLT or 3’ovr dsRNA, respectively) were mixed in annealing buffer (50 mM Tris pH 8.0, 20mM KCl). Mixtures were heated at 95°C for 2 min and allowed to cool to room temperature for 4 hrs. before purification after 8% native PAGE.

#### Nucleotides

ATP and ATP-γS were purchased from Sigma-Aldrich (St Louis, MO, USA) and used as received.

#### Sample preparation

Small-volume aliquots for fluorescence experiments were prepared immediately before use and contain 0.2 μM of Cy3 end-labeled dsRNA and, if used: 2 μM of dmDcr-2 protein (wild type or mutant), 2 μM of Loqs-PD, and 8 mM of ATP or ATP-γS. A full list of samples is provided in **Tables S1-S2**. Samples were allowed to equilibrate for 5 min. prior to time-resolved fluorescence measurements.

#### Time-resolved photoluminescence instrument, data collection and analysis

Low-background fluorescence measurements of chromophore-labeled dsRNA constructs were performed using a polarization-resolved ultrafast fluorescence instrument with few modifications from a previously described setup.(35) Instrumental and data analysis details in the Supporting Information.

## RESULTS AND DISCUSSION

The use of fluorescent probes as reporters of local conditions has been used extensively in biophysical studies, with a wide array of chromophores that serve as polarity, pH, viscosity, and ion concentration sensors.(36–44) Typical spectroscopic observables include brightness, spectral shifts in absorption/emission, fluorescence lifetime, and steady-state or time-resolved fluorescence anisotropy. In this study, we employ a combination of time-resolved photophysical observables of an indocarbocyanine dye (Cy3) to determine its intramolecular and intermolecular measures of rigidity. With these measurements, we characterize the local molecular dynamics at the binding site of dsRNA by dmDcr-2, a protein whose distinct domains offer selectivity toward BLT or 3’ovr substrates.

### Photophysics of a fluorescent label as a probe of local environment

The local environment surrounding a chromophore is coupled to its molecular dynamics and can strongly affect its photophysical behavior, which is the basis for fluorescence studies that probe biomolecular interactions using intrinsic or extrinsic chromophores.(45–58) Particularly, photoexcited indocarbocyanine dyes are known to undergo a cis-trans isomerization of their polymethine chain, which results in an efficient nonradiative decay pathway. When the local environment of indocarbocyanine dyes becomes more rigid, their excited state isomerization is hindered, which is reflected in an extension of their fluorescence lifetime and an increase in their brightness.(59–63) This effect has been observed in homogeneous bulk solutions in a viscosity- and temperature-dependent manner, and it is the basis of protein-induced fluorescence enhancement effects.(46) Beyond these excited state decay dynamics that reflect primarily an intramolecular process, collective processes can reflect important local environment properties – e.g., the three-dimensional reorientation of a group of chromophores can be observed in their time-resolved fluorescence anisotropy.(45, 47) A linearly polarized pump laser preferentially excites molecules whose transition dipole moments are aligned with its polarization, creating an anisotropic distribution of excited state dipoles even in an otherwise isotropic sample. If dye molecules are able to rotate in 3-dimensions, this initially-anisotropic excited state dipole distribution de-polarizes over a characteristic rotational diffusion time that depends strongly on the hydrodynamic radius of the rotating emitter. When rotation is unimpeded, the long-time average polarization is negligible. However, if the rotation of the emitters is hindered, a nonzero anisotropy can remain present at long times. This restricted motion is typically observed for dye-labeled nucleic acid-protein complexes, where the rotational diffusion of a chromophore also involves its immediate solvation shell, the macromolecule to which it is covalently attached, and nearby protein segments.(45) Here, we combine these intramolecular and collective measures of rigidity to characterize the local environment surrounding a fluorescent probe located at the terminus of dsRNA as it is recognized by dmDcr-2 as follows:

- to monitor the <u>intramolecular</u> rigidity of the fluorescent probe, we compute the mean fluorescence lifetime 〈τ〉 using the intensity-weighted photon arrival times (individual traces and details of their analysis given in **Figs. S1-S6** and **Tables S3-S4**), which we then normalize by the fluorescence lifetime of the free dye in solution, τ_Cy3_.
- the ability of the fluorescent probe to reorient via <u>collective</u> rotational diffusion processes is determined by the fraction of the initial anisotropy that remains at long times, *r*(∞)/*r*(0), obtained by nonlinear least-square fits to the anisotropy decays (individual traces and details of their analysis given in **Figs. S7-S12** and **Tables S5-S6**).

These complementary observations of intramolecular and collective processes provide a sensitive measure of the local molecular environment at the probe site (**Fig. 2**), and serve as a means to identify distinct modes of molecular recognition, determine the effect of partner proteins in overriding these molecular recognition pathways, and report on conformational changes of large biomolecular complexes as a result of ATP binding and hydrolysis.

**Figure 2.**
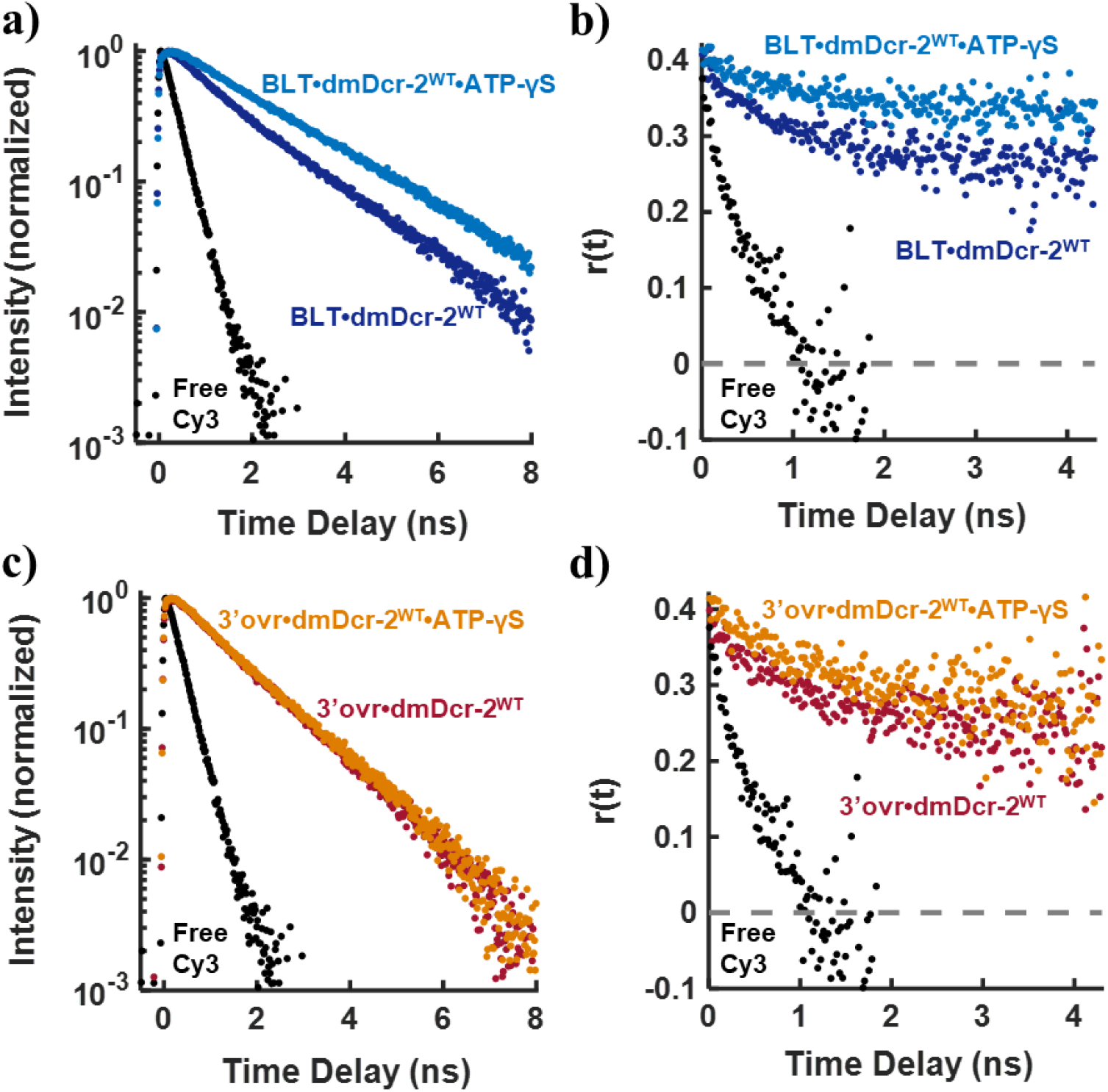
Simultaneous detection of time-resolved photoluminescence in orthogonal polarization channels is used to calculate **(a,c)** the rotationally-averaged fluorescence intensity, and **(b,d)** the time-resolved fluorescence anisotropy for Cy3 chromophores. Time-resolved photophysics of dyes covalently attached to dsRNA substrates (**a-b** BLT, **c-d** 3’ovr) display substantial differences that report on their local environment as a function of their molecular recognition by Dicer-2. For example, the local environment of BLT dsRNA bound by dmDcr-2^WT^ (dark blue) becomes increasingly rigid upon the addition of ATP-γS (light blue), but a much smaller difference is observed between the photophysics of Cy3 attached to 3’ovr dsRNA when bound by dmDcr-2^WT^ (red) vs. dmDcr-2^WT^•ATP-γS (orange). As a comparison, the fluorescence and transient anisotropy dynamics of free Cy3 in solution are also shown (black). A complete set of fluorescence and anisotropy traces are shown in **Fig. S1-S12**.

#### RNA-binding by distinct protein domains with substantial differences in local environment

Biochemical and structural evidence have led to a model for dmDcr-2 processing of dsRNA in which the helicase domain is essential for the binding and cleavage of BLT substrates, while the platform•PAZ domains play an important role in the processing of 3’ovr substrates – although overlap between binding sites has been detected in transient studies.(29) The helicase domain undergoes substantial conformational changes as a result of dsRNA and ATP binding, evidenced by the observation of protease-resistant fragments and differences in cryo-EM structures. These conformational changes result in the closing of the helicase pocket by a concerted motion of the Hel2 and Hel2i domains toward the Hel1 domain, sandwiching the dsRNA substrate.(24, 27) To determine whether the local environment at the dsRNA terminus can identify its mode of molecular recognition, we compared the intramolecular and collective measures of local rigidity – 〈τ〉/τ_Cy3_ and *r*(∞)/*r*(0), respectively – for end-labeled 52-basepair 3’ovr or BLT substrates as they are bound by wild type dmDcr-2 in the presence and absence of ATP-γS (**Fig. 3**).

**Figure 3.**
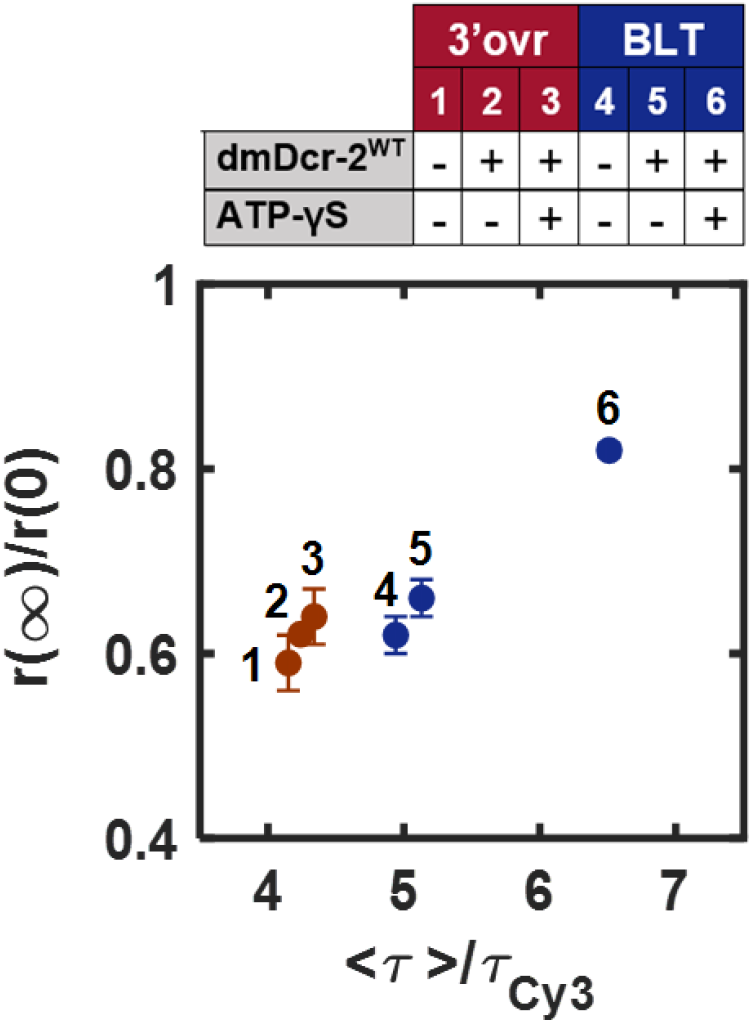
Combining intramolecular and collective measures of local rigidity enables the identification of dsRNA terminus-dependent binding environments of dmDcr-2. We calculate the normalized mean fluorescence lifetime and fractional residual anisotropy for each sample with (+) or without (−) dmDcr-2^WT^ and ATP-γS, as indicated. We observe that 3’ovr termini experience a minimal change in their local environment (markers 1-3 in red), while BLT termini experience a substantial increase in their local rigidity (markers 4-6 in blue).

Although not an optimal substrate, 3’ovr dsRNA can be cleaved by dmDcr-2^WT^ in the absence of ATP and regulatory proteins. However, the contribution of siRNA to our signal is not dominant given that 3’ovr recognition by dmDcr-2^WT^ and the non-cleaving mutant dmDcr-2^RIII^ show similar trends (**Fig. S13**), and because all complexes formed by 3’ovr and any dmDcr-2 variant belong to a single cluster of similar local environments (see Gaussian mixture model below).

The environment at the dsRNA terminus displays differences even in the absence of protein, with 3’ovr termini being less rigid than BLT termini (compare markers 1 and 4 in **Fig. 3**). This observation is consistent with the dependence of Cy3 emission on the terminal basepair it labels in DNA studies.(47) We observe marginal changes in the local environment experienced by 3’ovr dsRNA termini upon binding by dmDcr-2^WT^, and no further discernible change with the addition of ATP-γS (markers 1-2-3 in **Fig. 3**). In contrast, the local environment at BLT termini shows a small change upon binding by dmDcr-2^WT^, but a substantial rigidification when ATP-γS is added (markers 4-5-6 in **Fig. 3**). Comparing the progression in local environment at the dsRNA termini for 3’ovr and BLT substrates, it is clear that they reflect two very distinct modes of molecular recognition. We interpret these trends as the formation of an equilibrated complex between 3’ovr dsRNA and dmDcr-2^WT^ where the dsRNA terminus is recognized by the platform•PAZ domain and whose long-time equilibrium configuration is largely unaffected by conformational changes localized at the helicase domain. Conversely, dsRNA BLT termini are recognized by the helicase domain which provides a less flexible binding environment that is further rigidified by the ATP-γS-induced closing of the helicase pocket. Importantly, these results demonstrate the ability to differentiate between two substrate-specific modes of molecular recognition by the same protein using the information collected by a single fluorescent probe located at the binding site.

### Loqs-PD as a mediator of molecular recognition

The large differences in termini-dependent activity of dmDcr-2 are greatly reduced when the regulatory protein Loqs-PD is present. This effect relies on the ability of Loqs-PD to bind dsRNA in a termini-independent manner with two canonical dsRNA-binding motifs while also using its C-terminal FDF-like motif to interact with the Hel2 domain of dmDcr-2.(31) Models for the concerted biochemical function of Loqs-PD and dmDcr-2 include the co-localization of dsRNA substrates and Loqs-PD at the helicase domain of dmDcr-2 in a manner that enhances its activity toward both BLT and 3’ovr substrates and substantially reduces the differences in their cleavage rates, ATP-dependent behavior, and siRNA product size distribution. Here we interrogate whether the change in the biochemical activity of dmDcr-2 toward distinct dsRNA substrates is reflected in the molecular recognition of such substrates as sensed by a fluorescent probe at the termini.

While two distinct local binding environments for dsRNA were evident in the absence of Loqs-PD (**Fig. 3**), the dsRNA•dmDcr-2^WT^•Loqs-PD complexes show strikingly similar modes of molecular recognition independent of whether BLT or 3’ovr substrates are involved (compare trends of markers 1-2-3 and 4-5-6 **Fig. 4**). In both cases, a substantial rigidification of the local environment at the dsRNA terminus is observed upon binding to dmDcr-2^WT^•Loqs-PD, and a subsequent increase in the local rigidity occurs upon binding of ATP-γS. Both of these progressions in local binding environment are not only similar to one another, but they are also analogous to the molecular recognition of BLT dsRNA by dmDcr-2^WT^ in the absence of Loqs-PD (markers 4-5-6 **Fig. 3**). A similar effect is observed when using dmDcr-2^RIII^ or dmDcr-2^PP^ mutants (**Figs. S14-S15**), emphasizing that observed effects are related to the helicase domain and do not involve cleavage or the platform•PAZ domain. Importantly, we detect changes in both intramolecular (fluorescence lifetime) and collective (residual anisotropy) measures of rigidity at the dsRNA terminus. Thus, it is unlikely that these changes are simply reflective of a similar local environment with slower rotational diffusion due to the addition of another macromolecule to the biomolecular complex. What these data show is that Loqs-PD has a substantial effect on the mode of molecular recognition of dsRNA by dmDcr-2, reducing the differences in the binding environment for 3’ovr vs BLT substrates. Loqs-PD effectively overrides the way in which dmDcr-2^WT^ recognizes 3’ovr dsRNA substrates, while its effect on the mode of molecular recognition of BLT substrates by dmDcr-2^WT^ is not as pronounced. To provide a more complete description of the similarities and differences between the various local binding environments experienced by dsRNA as it is recognized by dmDcr-2, we include a larger set of observations and perform a cluster analysis that identifies distinct sub-populations that represent closely-related conformations of this large biomolecular complex.

**Figure 4.**
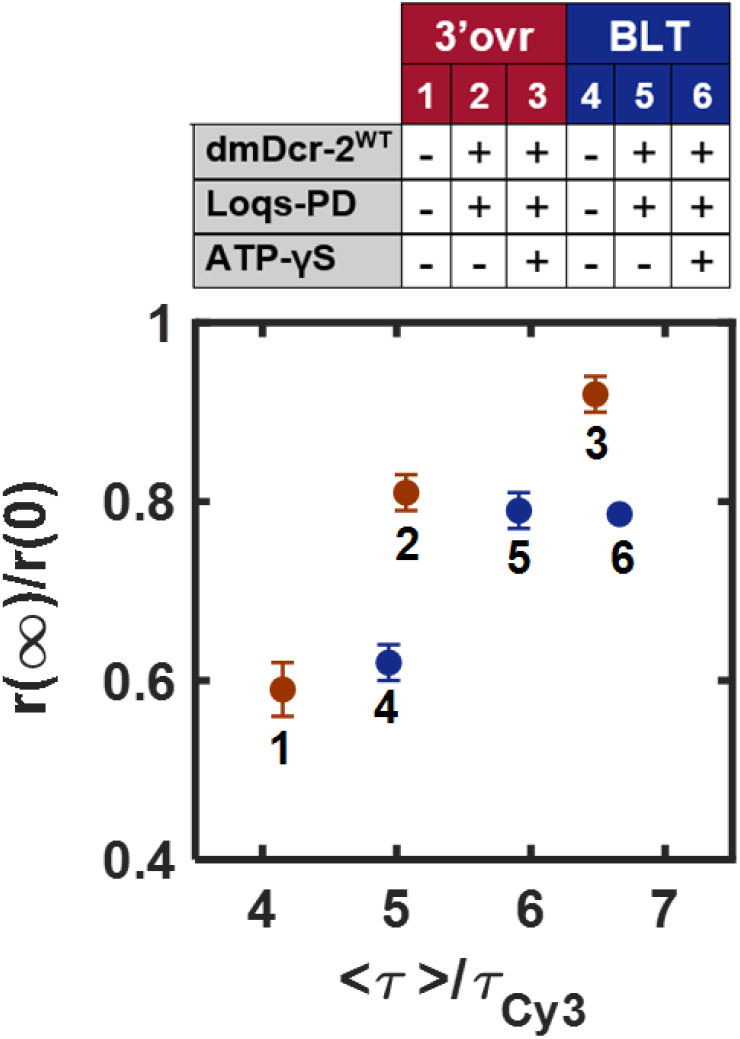
Intramolecular and collective measures of local rigidity sensed at dsRNA termini as they are bound by dmDcr-2^WT^ in the presence of the regulatory protein Loqs-PD. The progression in local environment sensed at 3’ovr (markers 1-3 in red) or BLT (markers 4-6 in blue) display a similar trend to one another, an indication that Loqs-PD overrides the termini-specific molecular recognition of dsRNA by dmDcr-2.

### Specific molecular recognition pathways lead to functional conformations

To identify differences and similarities in the local binding environment at the dsRNA terminus when recognized by dmDcr-2 as a function of nucleotide binding and hydrolysis, as well as its modulation by the regulatory protein Loqs-PD, we perform a cluster analysis using Gaussian mixture models. These cluster analyses (**Fig. 5**) describe a set of observations with a linear combination of Gaussian sub-populations and assign each element in the test set to one of such sub-populations (or clusters). We conduct this analysis for all BLT dsRNA samples in the presence of dmDcr-2^WT^ and several variants (dmDcr-2^ΔHel^, dmDcr-2^RIII^, and dmDcr-2^PP^), with and without ATP, ATP-γS, and Loqs-PD (**Fig. 5a**); an equivalent analysis was done for 3’ovr dsRNA samples (**Fig. 5b**).

**Figure 5.**
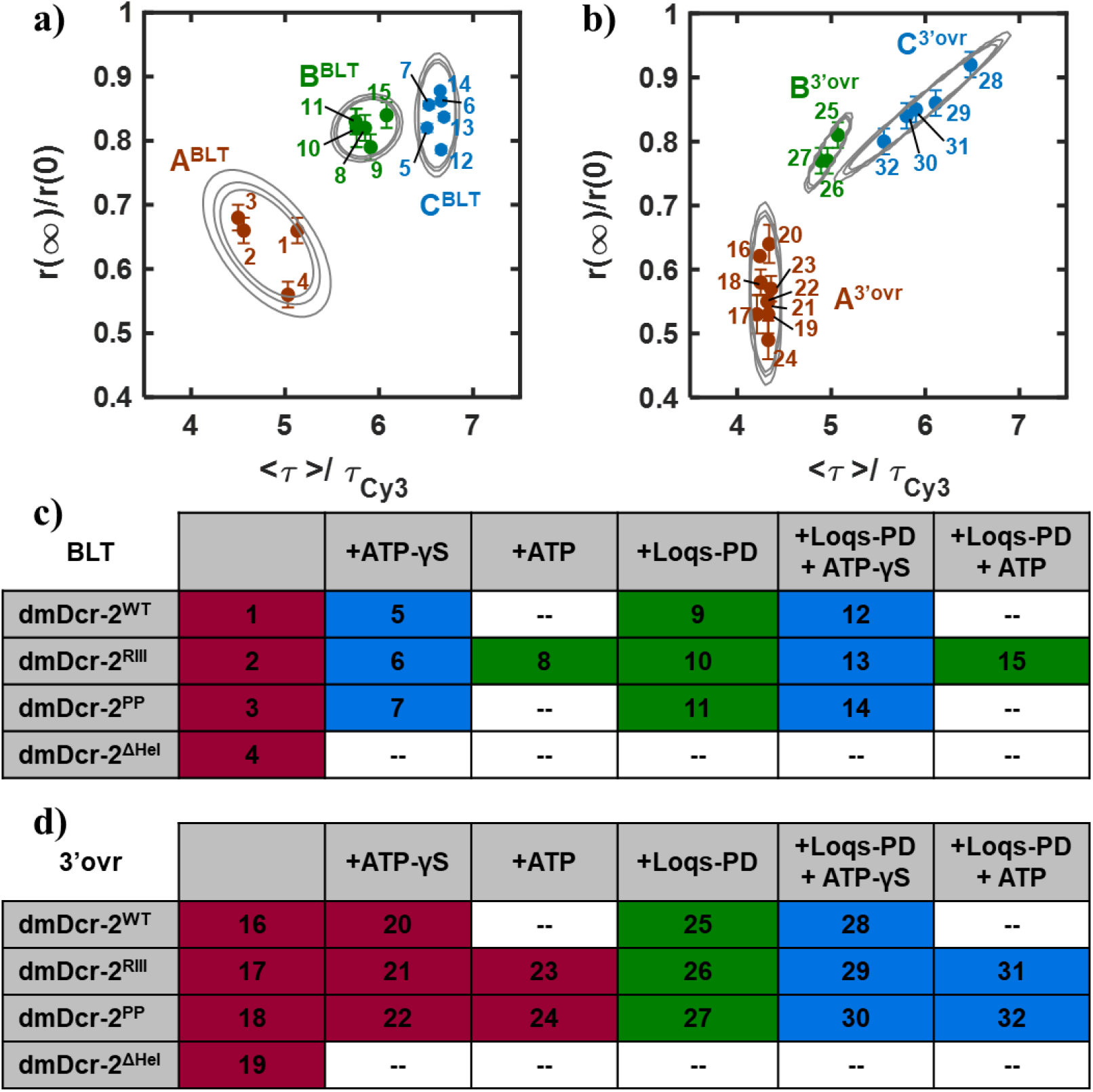
Gaussian mixture models for dsRNA•dmDcr-2 complexes for **(a)** BLT and **(b)** 3’ovr substrates. Three Gaussian sub-populations were observed in each case, and are labeled by cluster (A^BLT^-B^BLT^-C^BLT^ or A^3’ovr^-B^3’ovr^-C^3’ovr^) and color-coded to facilitate their distinction. These clusters correspond to distinct complex conformations with varying local environment at the dsRNA terminus. Samples are numbered as indicated in panels **c-d** (combinations not tested in this study are marked as double dashes). Model parameters for each cluster are listed in **Tables S7-S8**, and assignments of the complex conformational state for each cluster are discussed in the Main Text.

For both types of substrates, optimal fits are obtained when 3 Gaussian sub-populations are included, as described below (details in Supporting Information **Figs. S16-S17** and **Tables S7-S8**). As mentioned above, under equilibrium conditions dmDcr-2^WT^ and dmDcr-2^ΔHel^ can cleave 3’ovr dsRNA into siRNA, but the presence of siRNA is not a dominant factor in the local environment sensed at the dsRNA termini in these ensemble measurements at equilibrium. The Gaussian mixture models were constructed considering only complexes in which dmDcr-2^WT^ or a variant was present; however, a post-parametrization cluster assignment of samples that do not include dmDcr-2 (i.e., BLT or 3’ovr dsRNA with and without Loqs-PD) found them all to belong to the most flexible cluster (cluster A) in their corresponding Gaussian mixture models.

BLT dsRNA substrates experience a relatively flexible local environment at their terminus when bound by wild type and all variants of dmDcr-2 measured here, as long as no nucleotide or regulatory proteins are present (cluster A^BLT^, **Fig. 5a**). We interpret this cluster as the loosely bound conformations in the low-affinity BLT•dmDcr-2 complex. A much more rigid local environment at the BLT dsRNA terminus is observed upon binding by dmDcr-2^WT^ and all variants with a functional helicase domain in the presence of ATP-γS, irrespective of whether Loqs-PD is included or not (cluster C^BLT^, **Fig. 5a**). This cluster is thus interpreted to reflect conformations in which the BLT dsRNA terminus is located within a closed helicase domain but has not threaded through. The third type of environment reported by BLT dsRNA substrates (cluster B^BLT^, **Fig. 5a**) displays a slightly reduced rigidity from that of a closed helicase pocket. Importantly, this relative increase in flexibility is evidenced in the intramolecular dynamics and not on the collective, inter-molecular rotational diffusion. The complexes that belong to this cluster are those where BLT dsRNA interacts with a functional dmDcr-2 helicase in the presence of Loqs-PD, and those where ATP is present. As such, these complexes are interpreted to exist in a conformation where the terminus is either within an open helicase domain that is bound by Loqs-PD, or have threaded beyond a closed helicase pocket toward the platform•PAZ domain as a result of ATP hydrolysis.

The vast majority of 3’ovr dsRNA substrates form complexes with dmDcr-2 for which the local environment at the dsRNA terminus is flexible and shows little variation in intramolecular dynamics, but spans a considerable range in collective dynamics (cluster A^3’ovr^, **Fig. 5b**). This environment is experienced by 3’ovr dsRNA bound by dmDcr-2^WT^ and all dmDcr-2 variants measured in the absence of Loqs-PD, irrespective of the presence of ATP or ATP-γS. We therefore interpret the conformations in this cluster as an equilibrated bound state in which 3’ovr dsRNA interacts primarily with the platform•PAZ domain. While transient kinetic experiments have shown that 3’ovr dsRNA can also interact with the helicase domain in the presence of nucleotide,(29) these complexes have a shorter residence time than their BLT counterparts and are a minor component of the equilibrium conformations detected here. A second kind of local environment with intermediate rigidity is experienced by 3’ovr dsRNA termini bound by dmDcr-2^WT^ and variants with functional helicase domains in the presence of Loqs-PD (cluster B^3’ovr^, **Fig. 5b**). Given the considerable differences in the intramolecular and collective measures of local environment between the samples in cluster B^3’ovr^ and those in cluster A^3’ovr^, the conformations of 3’ovr•dmDcr-2•Loqs-PD are unlikely to reflect interaction of the 3’ovr terminus with the platform•PAZ domain. In turn, considering the similarities in the overall trends for molecular recognition that involves Loqs-PD to those for BLT dsRNA in the absence of Loqs-PD, it is possible that the complexes in cluster B^3’ovr^ reflect a Loqs-PD mediated interaction of 3’ovr dsRNA with an open helicase pocket. Lastly, the most rigid local environments experienced by 3’ovr termini are those when they are bound by a functional dmDcr-2 helicase with the assistance of Loqs-PD in the presence of ATP or ATP-γS (cluster C^3’ovr^, **Fig. 5b**). We do not observe separate clusters for complexes including ATP vs. ATP-γS as it was the case for BLT dsRNA. However, pairwise comparisons for 3’ovr•dmDcr-2•Loqs-PD•nucleotide complexes with the same dmDcr-2 variant (**Fig. 5b**, points 29 vs. 31 for dmDcr-2^RIII^ and 30 vs. 32 for dmDcr-2^PP^) show that the environment at the 3’ovr terminus is more rigid in the absence of hydrolysis in each case, as expected. All clusters extracted for 3’ovr substrates are less rigid than their BLT counterparts (most notably A^BLT^ vs. A^3’ovr^ and B^BLT^ vs. B^3’ovr^), and they also display substantial variability in the local binding environments sensed at 3’ovr termini (consider spread in clusters A^3’ovr^ and C^3’ovr^). These less rigid and more variable environments at 3’ovr termini can be related to a combination of several factors: an increased local flexibility at 3’ovr termini due to unpaired nucleotides, a decrease in the efficiency with which the helicase domain clamps onto 3’ovr vs. BLT dsRNA substrates (as suggested by a lower amount of protease resistant fragment in limited proteolysis studies and the smaller component of heterogeneous siRNA products from 3’ovr dsRNA), or to variability in the location of the dsRNA terminus as it threads from the helicase to the platform•PAZ end of the protein.

Beyond interpreting conformations at the individual cluster level, we aim to connect these local binding environment observations to the biochemical activity of dmDcr-2 in terms of its molecular recognition properties. We consider the formation of the biomolecular complex in the absence (***case 1***) or presence of Loqs-PD (***case 2***), and label the states involved in the formation of the biomolecular complex (**I** through **IV**):

***Case 1:***

**(I)** dsRNA ⟶ **(II)** dsRNA•dmDcr-2 ⟶ **(III)** dsRNA•dmDcr-2•ATP-γS

***Case 2:***

**(I)** dsRNA ⟶ **(II)** dsRNA•Loqs-PD ⟶ **(III)** dsRNA•Loqs-PD•dmDcr-2 ⟶ **(IV)** dsRNA•Loqs-PD•dmDcr-2•ATP-γS and employ the cluster analysis as a proxy for the local conformations experienced by BLT or 3’ovr dsRNA termini. We restrict these comparisons to dmDcr-2 proteins with functional helicase domains (WT, RIII, or PP). For states **I**-**II**-**III**shown in ***case 1*** (without Loqs-PD), BLT substrates are found in distinct clusters (A^BLT^-A^BLT^-C^BLT^) while 3’ovr substrates remain in the same cluster (A^3’ovr^-A^3’ovr^-A^3’ovr^). For states **I-II-III-IV** shown in ***case 2*** (with Loqs-PD), both types of dsRNA substrates are found in states that belong to distinct clusters (A^BLT^-A^BLT^-B^BLT^-C^BLT^, and A^3’ovr^-A^3’ovr^-B^3’ovr^-C^3’ovr^). While the specifics of the conformation in each cluster differ (e.g., B^BLT^ vs B^3’ovr^), we draw attention to how large changes in local binding environment at the dsRNA termini are reflected by a switch in cluster assignment, while smaller differences in local binding environment mean that two samples are assigned to the same cluster. Therefore, we identify three distinct modes of molecular recognition reflected on the equilibrium, ensemble-averaged configurations at the dsRNA terminus: (1) strong, nucleotide-dependent BLT binding by the helicase domain, (2) weaker, nucleotide-independent 3’ovr binding by the platform•PAZ domain, and (3) strong, Loqs-PD mediated, and nucleotide-dependent binding of BLT or 3’ovr by the helicase domain. These substrate-selective (case 1) or substrate-independent (case 2) molecular recognition properties of dmDcr-2 are strongly correlated to the differences in biochemical activity of dmDcr-2 toward BLT and 3’ovr dsRNA, as well as the substantial reduction of such differences in the presence of Loqs-PD.

## CONCLUSION

Advancing our understanding of the role played by local binding environment in the process of molecular recognition between large biomolecules requires new, molecular-scale observations in well-controlled interactions. To probe differences in local binding environment for the terminus-specific molecular recognition of dsRNA by dmDcr-2, we employ the photophysical behavior of a chromophore placed at the dsRNA terminus. Using a cluster analysis based solely on the intramolecular and collective photophysical observables of these fluorescent probes, it is possible to identify the local binding environments reflective of distinct conformations of the biomolecular complex with strong correlation to biochemical observables such as cleavage rate, ATP dependence, and product size distribution. We detect a clear differentiation of multiple conformations in a substrate-specific manner and their dependence on regulatory factors, and we connect these observations to distinct modes of molecular recognition that involve multiple protein domains. While the helicase domain offers a rigid binding environment for dsRNA, a less rigid binding environment is experienced by dsRNA at the platform•PAZ domain. Binding of ATP-γS at the helicase domain leads to conformational changes that substantially rigidify the local environment at the dsRNA terminus whenever it is located within the helicase pocket. Moreover, ATP hydrolysis has been shown to be essential for the processing of substrates recognized by the helicase domain, and its effect on the local environment at the dsRNA terminus is a noticeable reduction of rigidity – in agreement with the expectation that ATP hydrolysis is related to dsRNA threading through the helicase pocket and toward the RNase III cleaving site and further to the platform•PAZ domain. Previously reported changes in biochemical activity toward suboptimal 3’ovr dsRNA substrates mediated by the RNA-binding protein Loqs-PD are shown to be due to the modification of its molecular recognition pathway. More generally, these studies of local binding environment in biomolecular complexes offer an opportunity to disentangle the connection between molecular recognition and biochemical function in large proteins whose multiple domains can interact with a broad diversity of substrates. This molecular-scale characterization using dynamic spectroscopic observables represents a generalizable approach to enable detailed studies into the diversity of molecular recognition modes employed by RNA-binding proteins.

## ACKNOWLEDGMENTS

This work was supported by funds to B.L.B. from the National Institute of General Medical Sciences (R01GM121706) and startup funds to R.N. from the University of Utah Chemistry Department.

## Author Contributions

RKS, BLB and RN designed experiments. MJ and RKS performed the experiments. RKS prepared dsRNA and proteins. MJ and RN analyzed data and wrote the manuscript with input from all authors.

## REFERENCES

1. Castello, A., B. Fischer, M.W. Hentze, and T. Preiss. 2013. RNA-binding proteins in Mendelian disease. Trends in Genetics. 29:318–327.

2. Hentze, M.W., A. Castello, T. Schwarzl, and T. Preiss. 2018. A brave new world of RNA-binding proteins. Nat Rev Mol Cell Biol. 19:327–341.

3. Lunde, B.M., C. Moore, and G. Varani. 2007. RNA-binding proteins: modular design for efficient function. Nature Reviews Molecular Cell Biology. 8:479–490.

4. Lenzken, S.C., T. Achsel, M.T. Carrì, and S.M.L. Barabino. 2014. Neuronal RNA-binding proteins in health and disease. WIREs RNA. 5:565–576.

5. Nogales, E. 2016. The development of cryo-EM into a mainstream structural biology technique. Nature Methods. 13:24–27.

6. Fernandez-Leiro, R., and S.H.W. Scheres. 2016. Unravelling biological macromolecules with cryo-electron microscopy. Nature. 537:339–346.

7. Schlichting, I., and J. Miao. 2012. Emerging opportunities in structural biology with X-ray free-electron lasers. Current Opinion in Structural Biology. 22:613–626.

8. Shi, Y. 2014. A Glimpse of Structural Biology through X-Ray Crystallography. Cell. 159:995–1014.

9. Hennig, J., and M. Sattler. 2014. The dynamic duo: Combining NMR and small angle scattering in structural biology. Protein Science. 23:669–682.

10. 2014. Present and future of NMR for RNA–protein complexes: A perspective of integrated structural biology. Journal of Magnetic Resonance. 241:126–136.

11. Daubner, G.M., A. Cléry, and F.H.-T. Allain. 2013. RRM–RNA recognition: NMR or crystallography…and new findings. Current Opinion in Structural Biology. 23:100–108.

12. Ke, A., and J.A. Doudna. 2004. Crystallization of RNA and RNA–protein complexes. Methods. 34:408–414.

13. Cheatham, T.E. 2004. Simulation and modeling of nucleic acid structure, dynamics and interactions. Current Opinion in Structural Biology. 14:360–367.

14. McCammon, J.A. 1998. Theory of biomolecular recognition. Current Opinion in Structural Biology. 8:245–249.

15. Boehr, D.D., R. Nussinov, and P.E. Wright. 2009. The role of dynamic conformational ensembles in biomolecular recognition. Nature Chemical Biology. 5:789–796.

16. Fox, J.M., M. Zhao, M.J. Fink, K. Kang, and G.M. Whitesides. 2018. The Molecular Origin of Enthalpy/Entropy Compensation in Biomolecular Recognition. Annual Review of Biophysics. 47:223–250.

17. Mackereth, C.D., and M. Sattler. 2012. Dynamics in multi-domain protein recognition of RNA. Current Opinion in Structural Biology. 22:287–296.

18. Henzler-Wildman, K., and D. Kern. 2007. Dynamic personalities of proteins. Nature. 450:964–972.

19. López, C.J., S. Oga, and W.L. Hubbell. 2012. Mapping Molecular Flexibility of Proteins with Site-Directed Spin Labeling: A Case Study of Myoglobin. Biochemistry. 51:6568–6583.

20. Hartman, E., Z. Wang, Q. Zhang, K. Roy, G. Chanfreau, and J. Feigon. 2013. Intrinsic Dynamics of an Extended Hydrophobic Core in the S. cerevisiae RNase III dsRBD Contributes to Recognition of Specific RNA Binding Sites. Journal of Molecular Biology. 425:546–562.

21. Frauenfelder, H., S.G. Sligar, and P.G. Wolynes. 1991. The energy landscapes and motions of proteins. Science. 254:1598–1603.

22. Mackereth, C.D., T. Madl, S. Bonnal, B. Simon, K. Zanier, A. Gasch, V. Rybin, J. Valcárcel, and M. Sattler. 2011. Multi-domain conformational selection underlies pre-mRNA splicing regulation by U2AF. Nature. 475:408–411.

23. Tants, J.-N., S. Fesser, T. Kern, R. Stehle, A. Geerlof, C. Wunderlich, M. Juen, C. Hartlmüller, R. Böttcher, S. Kunzelmann, O. Lange, C. Kreutz, K. Förstemann, and M. Sattler. 2017. Molecular basis for asymmetry sensing of siRNAs by the Drosophila Loqs-PD/Dcr-2 complex in RNA interference. Nucleic Acids Res. 45:12536–12550.

24. Sinha, N.K., J. Iwasa, P.S. Shen, and B.L. Bass. 2018. Dicer uses distinct modules for recognizing dsRNA termini. Science. 359:329–334.

25. Lee, Y.S., K. Nakahara, J.W. Pham, K. Kim, Z. He, E.J. Sontheimer, and R.W. Carthew. 2004. Distinct Roles for Drosophila Dicer-1 and Dicer-2 in the siRNA/miRNA Silencing Pathways. Cell. 117:69–81.

26. Mussabekova, A., L. Daeffler, and J.-L. Imler. 2017. Innate and intrinsic antiviral immunity in Drosophila. Cell. Mol. Life Sci. 74:2039–2054.

27. Sinha, N.K., K.D. Trettin, P.J. Aruscavage, and B.L. Bass. 2015. Drosophila Dicer-2 Cleavage Is Mediated by Helicase- and dsRNA Termini-Dependent States that Are Modulated by Loquacious-PD. Molecular Cell. 58:406–417.

28. Donelick, H.M., L. Talide, M. Bellet, J. Aruscavage, E. Lauret, E. Aguiar, J.T. Marques, C. Meignin, and B.L. Bass. 2020. In vitro studies provide insight into effects of Dicer-2 helicase mutations in Drosophila melanogaster. RNA. rna.077289.120.

29. Singh, R.K., M. Jonely, E. Leslie, N.A. Rejali, R. Noriega, and B.L. Bass. 2020. A real-time, transient kinetic study of Drosophila melanogaster Dicer-2 elucidates mechanism of termini-dependent cleavage of dsRNA. bioRxiv. 2020.09.29.319475.

30. Jankowsky, E. 2011. RNA helicases at work: binding and rearranging. Trends in Biochemical Sciences. 36:19–29.

31. Trettin, K.D., N.K. Sinha, D.M. Eckert, S.E. Apple, and B.L. Bass. 2017. Loquacious-PD facilitates Drosophila Dicer-2 cleavage through interactions with the helicase domain and dsRNA. PNAS. 114:E7939–E7948.

32. Ahmad, S., and S. Hur. 2015. Helicases in Antiviral Immunity: Dual Properties as Sensors and Effectors. Trends in Biochemical Sciences. 40:576–585.

33. Hansen, S.R., A.M. Aderounmu, H.M. Donelick, and B.L. Bass. 2019. Dicer’s Helicase Domain: A Meeting Place for Regulatory Proteins. Cold Spring Harb Symp Quant Biol. 84:185–193.

34. Kandasamy, S.K., and R. Fukunaga. 2016. Phosphate-binding pocket in Dicer-2 PAZ domain for high-fidelity siRNA production. PNAS. 113:14031–14036.

35. Jonely, M., and R. Noriega. 2020. Role of Polar Protic Solvents in the Dissociation and Reactivity of Photogenerated Radical Ion Pairs. J. Phys. Chem. B. 124:3083–3089.

36. Lakowicz, J.R. 2006. Principles of Fluorescence Spectroscopy. Third Ed. Springer.

37. Mandal, P.K., and A. Samanta. 2005. Fluorescence Studies in a Pyrrolidinium Ionic Liquid: Polarity of the Medium and Solvation Dynamics. J. Phys. Chem. B. 109:15172–15177.

38. Das, R., D. Guha, S. Mitra, S. Kar, S. Lahiri, and S. Mukherjee. 1997. Intramolecular Charge Transfer as Probing Reaction: Fluorescence Monitoring of Protein−Surfactant Interaction. J. Phys. Chem. A. 101:4042–4047.

39. Zhujun, Z., and W.R. Seitz. 1984. A fluorescence sensor for quantifying pH in the range from 6.5 to 8.5. Analytica Chimica Acta. 160:47–55.

40. Saari, L.A., and W.R. Seitz. 1982. pH sensor based on immobilized fluoresceinamine. Anal. Chem. 54:821–823.

41. Peng, X., Z. Yang, J. Wang, J. Fan, Y. He, F. Song, B. Wang, S. Sun, J. Qu, J. Qi, and M. Yan. 2011. Fluorescence Ratiometry and Fluorescence Lifetime Imaging: Using a Single Molecular Sensor for Dual Mode Imaging of Cellular Viscosity. J. Am. Chem. Soc. 133:6626–6635.

42. Haidekker, M.A., T.P. Brady, D. Lichlyter, and E.A. Theodorakis. 2006. A Ratiometric Fluorescent Viscosity Sensor. J. Am. Chem. Soc. 128:398–399.

43. Lin, H.-Y., P.-Y. Cheng, C.-F. Wan, and A.-T. Wu. 2012. A turn-on and reversible fluorescence sensor for zinc ion. Analyst. 137:4415–4417.

44. Crivat, G., K. Kikuchi, T. Nagano, T. Priel, M. Hershfinkel, I. Sekler, N. Rosenzweig, and Z. Rosenzweig. 2006. Fluorescence-Based Zinc Ion Sensor for Zinc Ion Release from Pancreatic Cells. Anal. Chem. 78:5799–5804.

45. Sanborn, M.E., B.K. Connolly, K. Gurunathan, and M. Levitus. 2007. Fluorescence Properties and Photophysics of the Sulfoindocyanine Cy3 Linked Covalently to DNA. J. Phys. Chem. B. 111:11064–11074.

46. Stennett, E.M.S., M.A. Ciuba, S. Lin, and M. Levitus. 2015. Demystifying PIFE: The Photophysics Behind the Protein-Induced Fluorescence Enhancement Phenomenon in Cy3. J. Phys. Chem. Lett. 6:1819–1823.

47. Spiriti, J., J.K. Binder, M. Levitus, and A. van der Vaart. 2011. Cy3-DNA Stacking Interactions Strongly Depend on the Identity of the Terminal Basepair. Biophysical Journal. 100:1049–1057.

48. Fürstenberg, A., M.D. Julliard, T.G. Deligeorgiev, N.I. Gadjev, A.A. Vasilev, and E. Vauthey. 2006. Ultrafast Excited-State Dynamics of DNA Fluorescent Intercalators: New Insight into the Fluorescence Enhancement Mechanism. J. Am. Chem. Soc. 128:7661–7669.

49. Remington, J.M., A.M. Philip, M. Hariharan, and B. Kohler. 2016. On the origin of multiexponential fluorescence decays from 2-aminopurine-labeled dinucleotides. J. Chem. Phys. 145:155101.

50. Jean, J.M., and K.B. Hall. 2001. 2-Aminopurine fluorescence quenching and lifetimes: Role of base stacking. PNAS. 98:37–41.

51. Jean, J.M., and K.B. Hall. 2002. 2-Aminopurine Electronic Structure and Fluorescence Properties in DNA. Biochemistry. 41:13152–13161.

52. Nguyen, B., M.A. Ciuba, A.G. Kozlov, M. Levitus, and T.M. Lohman. 2019. Protein Environment and DNA Orientation Affect Protein-Induced Cy3 Fluorescence Enhancement. Biophysical Journal. 117:66–73.

53. Rajendran, S., M.J. Jezewska, and W. Bujalowski. 2001. Multiple-Step Kinetic Mechanisms of the ssDNA Recognition Process by Human Polymerase β in Its Different ssDNA Binding Modes. Biochemistry. 40:11794–11810.

54. Luo, G., M. Wang, W.H. Konigsberg, and X.S. Xie. 2007. Single-molecule and ensemble fluorescence assays for a functionally important conformational change in T7 DNA polymerase. PNAS. 104:12610–12615.

55. Bjornson, K.P., K.J.M. Moore, and T.M. Lohman. 1996. Kinetic Mechanism of DNA Binding and DNA-Induced Dimerization of the Escherichia coli Rep Helicase. Biochemistry. 35:2268–2282.

56. Overman, L.B., W. Bujalowski, and T.M. Lohman. 1988. Equilibrium binding of Escherichia coli single-strand binding protein to single-stranded nucleic acids in the (SSB)65 binding mode. Cation and anion effects and polynucleotide specificity. Biochemistry. 27:456–471.

57. Fischer, C.J., N.K. Maluf, and T.M. Lohman. 2004. Mechanism of ATP-dependent Translocation of E.coli UvrD Monomers Along Single-stranded DNA. Journal of Molecular Biology. 344:1287–1309.

58. Kozlov, A.G., R. Galletto, and T.M. Lohman. 2012. SSB–DNA Binding Monitored by Fluorescence Intensity and Anisotropy. In: Keck JL, editor. Single-Stranded DNA Binding Proteins: Methods and Protocols. Totowa, NJ: Humana Press. pp. 55–83.

59. Åkesson, E., V. Sundström, and T. Gillbro. 1985. Solvent-dependent barrier heights of excited-state photoisomerization reactions. Chemical Physics Letters. 121:513–522.

60. Widengren, J., and P. Schwille. 2000. Characterization of Photoinduced Isomerization and Back-Isomerization of the Cyanine Dye Cy5 by Fluorescence Correlation Spectroscopy. J. Phys. Chem. A. 104:6416–6428.

61. Jia, K., Y. Wan, A. Xia, S. Li, F. Gong, and G. Yang. 2007. Characterization of Photoinduced Isomerization and Intersystem Crossing of the Cyanine Dye Cy3. J. Phys. Chem. A. 111:1593–1597.

62. Stennett, E.M.S., N. Ma, A. van der Vaart, and M. Levitus. 2014. Photophysical and Dynamical Properties of Doubly Linked Cy3–DNA Constructs. J. Phys. Chem. B. 118:152–163.

63. Hart, S.M., J.L. Banal, M. Bathe, and G.S. Schlau-Cohen. 2020. Identification of Nonradiative Decay Pathways in Cy3. J. Phys. Chem. Lett. 11:5000–5007.

